# Deep learning based on multi-omics integration identifies potential therapeutic targets in breast cancer

**DOI:** 10.1101/2022.01.18.476842

**Authors:** Xingxin Pan, Brandon Burgman, Nidhi Sahni, S. Stephen Yi

**Affiliations:** Department of Oncology, Livestrong Cancer Institutes, Dell Medical School, The University of Texas at Austin, Austin, TX 78712, USA; Interdisciplinary Life Sciences Graduate Programs (ILSGP), College of Natural Sciences, The University of Texas at Austin, Austin, TX 78712, USA; Department of Epigenetics and Molecular Carcinogenesis, The University of Texas MD Anderson Cancer Center, Smithville, TX 78957, USA; Department of Bioinformatics and Computational Biology, The University of Texas MD Anderson Cancer Center, Houston, TX 77030, USA; Quantitative and Computational Biosciences Program, Baylor College of Medicine, Houston, TX 77030, USA; Oden Institute for Computational Engineering and Sciences (ICES), The University of Texas at Austin, Austin, TX 78712, USA; Department of Biomedical Engineering, Cockrell School of Engineering, The University of Texas at Austin, Austin, TX 78712, USA

**Keywords:** multi-omics, deep learning model, data integration, patient stratification, breast cancer

## Abstract

Effective and precise classification of breast cancer patients for their disease risks is critical to improve early diagnosis and patient survival. In the recent past, a significant amount of multi-omics data derived from cancer patients has emerged. However, a robust framework for integrating multi-omics data to subgroup cancer patients and predict survival prognosis is still lacking. In addition, effective therapeutic targets for treating breast cancer patients with poor prognoses are in dire need. To begin to resolve this difficulty, we developed and optimized a sophisticated deep learning-based model in breast cancer that can accurately stratify patients based on their prognosis. We built a survival-associated predictive framework integrating transcription profile, miRNA expression, somatic mutations, copy number variation, DNA methylation and protein expression. This framework achieved promising performance in distinguishing high-risk breast cancer patients from those with good prognoses. Furthermore, we constructed multiple fully connected neural networks that are trained on prioritized multi-omics signatures or even only potential single-omics signatures, based on our customized scoring system. Together, the landmark multi-omics signatures we identified may serve as potential therapeutic targets in breast cancer.

## Introduction

Breast cancer is a leading cancer type responsible for mortality in women worldwide, and each year, more than 300,000 women in the United States are diagnosed with breast cancer [1, 2]. Breast cancer incidence is increasing in almost all ethnicities in the world and this cancer has been the second most common cause of cancer death among women in the world [1]. Although overall death rates from breast cancer have decreased in the past 20 years, the high level of heterogeneity in breast cancer along with the complex biological factors makes the prognosis prediction very challenging [1, 3]. For example, triple-negative breast cancer is thought to be more aggressive and has a poorer prognosis than other subtypes of breast cancer like Luminal A [3, 4]. Moreover, treatment strategies in these high-risk breast cancers are still limited, imposing urgent needs for developing computational methods to distinguish high-risk breast cancer patients from those low-risk patients and discover novel therapeutic targets effectively [4–6].

To understand the heterogeneity among breast cancer, quite a few works have been done to identify breast cancer molecular subtypes [7–11]. Multiple novel molecular subtypes are identified, ranging from 2 to 5, based on various data like histopathology and gene expression profiles [7–12]. However, most of these works explore the molecular subtypes without taking survival prognosis into account, making the identified subtypes less valuable in clinical practice [7–13]. Instead, survival time is a success factor when classifying breast cancer patients into valuable subtypes with differential survival prognoses. Moreover, it remains challenging in discovering effective therapeutic targets for breast cancer especially high-risk breast cancer subtypes in the community. Therefore, computational methods to robustly classify breast cancer patients into survival-associated subtypes and to discover potential therapeutic targets are urgently needed.

With the advance of artificial intelligence and high-performance computing, machine learning technologies especially deep neural networks are playing a more and more active role in computational biology and gaining significant successes in various fields [14–18]. For example, deep learning has been applied to find gene expression-associated genetic variants, DNA methylation, biological image analysis, leading to great breakthroughs in these fields [14–18]. The application of deep learning to omics data is a promising area that is revolutionizing genome research.

To address the outstanding issues in breast cancer, for the first time, we utilized deep learning computational framework on a comprehensive multi-omics dataset in breast cancer, including mRNA profile, miRNA expression, DNA methylation, somatic mutation, RPPA protein expression, and copy number variation profiles. Considering multi-omics are high-dimensional features, we adopted and optimized one autoencoder framework to remodel multi-omics data using complex functions in the network and extracted low-dimensional informative features from the model [19, 20]. We classified breast cancer patients into two subgroups with significant differences in survival prognosis based on survival-associated low-dimensional features from the bottleneck in the autoencoder [19, 20]. These two subgroups represent two distinguished survival prognosis states in breast cancer patient populations, namely high-risk subgroup and low-risk subgroup. After extracting potential multi-omics features associated with breast cancer subgroups, we selected the top ranked multi-omics features as lardmark signatures. These potential signatures empower us to better understand breast cancer development and progression from the multi-omics aspect. Furthermore, we trained and optimized multiple fully connected neural networks, and these specified networks are proved to be robust and effective in subgrouping breast cancer patients compared with other benchmark prediction models based on multi-omics signatures or single-type omics signatures. Lastly, given that fully connected neural networks achieve robust predictions in subgrouping breast cancer patients, we evaluated and ranked all potential multi-omics signatures based on the intrinsic information in the neural networks, and some of the landmark signatures we prioritized may serve as putative therapeutic targets in breast cancer.

## Methods

### Datasets employed in this study

In this study, we used a comprehensive TCGA BRCA cohort. We obtained multi-omics breast cancer datasets, including RNA sequencing data (TPM normalized gene expression quantification), protein expression data (Reverse Phase Protein Array RPPA), miRNA-seq expression data (reads per million for miRNA mapping to miRbase 20), DNA methylation data (Infinium HumanMethylation450 BeadChip), copy number variation data (Affymetrix SNP Array 6.0) and somatic mutation data (DNA sequencing). For copy number variation data, we calculated a gene-level copy number value as the average copy number of the genomic region of a gene; for the DNA methylation, we mapped CpG islands within 1500 base pairs (bp) ahead of transcriptional start sites (TSS) and averaged their values as the methylation value of a gene; for somatic mutation data, we transformed the data into a matrix, where each row is one specific mutation of a gene, each column is a patient sample pair and each cell includes 0/1 elements, where 1 indicates the patient sample pair has this mutation while 0 indicates not. The data processed above and clinical patient information download were implemented by TCGA-Assembler 2 [21]. When it comes to the missing values, two steps were performed: we removed the samples where more than 20% of features are missing and we removed the features that have zero value in more than 20% of patients; we filled out the missing expression values via impute package [22, 23].

### Generation of low-dimensional transformed features using a deep learning framework

We treated each type of omics data such as RNA-seq as a matrix, where rows represent samples and each column represents the expression of each gene. The same data preprocessing could be applied to the other types of omics, including somatic mutation profiles where each column represents one specific mutation of each gene, protein expression profiles where each column represents the expression of each protein, copy number variation where each column represents CNV of each gene, DNA methylation profiles where each column represents methylation expression of each gene, and miRNA profiles where each column represents the expression of each miRNA. We stacked these matrices and normalized them to form one integrated matrix which could represent high-dimensional multi-omics features for each patient. After that, we took the integrated matrices as input in the autoencoder.

An autoencoder is one type of artificial neural network and it is proved to learn efficient representation from a mass of features in an unsupervised manner and achieve dimensionality reduction effectively [19, 20]. Here, we took advantage of the autoencoder to transform high-dimensional multi-omics information to low-dimensional features.

### The architecture of the autoencoder

We developed one autoencoder for automatic feature extraction. Briefly speaking, being normalized and scaled, the integrated multi-omics matrix is taken as an input. In the autoencoder, several hyperparameters and parameters needed to be optimized for the best fit during the training process, including the number of neural net layers, dropout rate and the number of neurons of each layer, batch size, etc. The best hyperparameter and parameter combination was validated via k-fold cross-validation. To get the best combination of hyperparameters in the high-dimensional hyperparameter space effectively, Bayesian optimization was adopted in the training process. Finally, the best autoencoder was obtained based on the integrated matrix data. The optimized autoencoder consists of three dense neural net layers (1000, 100, and 1000 nodes separately) with two BatchNormalization and dropout layers placed between the dense neural net layers [24, 25]. Activation function was tanh in those densely connected layers, and for obtaining better local minima of parameters, adam was adopted as gradient descent optimizer [26]. The loss function of the autoencoder was binary cross-entropy and the autoencoder was trained for a maximum of 300 epochs. Besides, as an effective regularization method, early stopping was also adopted to terminate training with specific patience once the model performance was no longer improved on the validation dataset to avoid overfitting [27].

After the autoencoder was finalized, we made use of the trained encoder of the autoencoder based on the input omics data and obtained 100 transformed low-dimensional features from the bottleneck layer to represent multi-omics features for each patient.

### Transformed feature selection and clustering

The autoencoder reduced the initial high-dimensional information to 100 low-dimensional features obtained from the bottleneck layer. Next for each of these transformed features produced by the optimized autoencoder, we combined clinical information and built a univariate Cox-PH model for each feature, and then selected informative features from which a significant Cox-PH model was obtained (both Wald and Log-rank P-value < 0.05). We further used these informative features to cluster the samples using partitioning around medoids, a more robust version of K-means [28]. To determine the optimal number of clusters, we calculated two metrics from multiple testing numbers of clusters, including Silhouette index and Calinski-Harabasz criterion.

### Molecular signatures for breast cancer subgrouping

After obtaining the labels from partitioning around medoids, we identified the potential multi-omics features that are most correlated with the risk subgroup of patients via Wilcoxon. We obtained 2,122 potential CNV features, 12,658 potential gene expression features, 6,419 potential methylation features, 38 potential protein expression features, 121 potential somatic mutation features, and 417 potential miRNA expression features. We selected top potential features according to P-values as potential multi-omics signatures, including 424 CNV signatures, 1,265 gene expression features, 641 methylation features, 38 protein expression signatures, 121 somatic mutation signatures, and 417 miRNA signatures.

### Subgroup prediction via deep neural networks

After getting potential multi-omics signatures, we sought to establish a robust and effective prediction model that could achieve breast cancer patient stratification well only based on the potential multi-omics signatures or single-omics signatures. More specifically, to predict subgroups of breast cancer patients based on multi-omics data, we built and optimized subgrouping prediction models from the combination of potential multi-omics signatures. In addition, we devised and optimized subgrouping prediction models from each single-omics data, using the corresponding potential signatures respectively. To guarantee the robust classification of breast cancer patients, we constructed multiple prediction models, including fully connected neural networks (FCN), support vector machine (SVM), random forest (RF), LogitBoost (LGB), and Naive Bayes (NB). We constructed multiple parameter spaces for each prediction model and carried out grid search to tune model parameters to find the best model with optimal parameters.

### Data partitioning and processing

We randomly split 70% of the breast cancer samples as training sets and 30% of the breast cancer samples as testing sets to guarantee a reliable performance evaluation. We determined the known labels of the breast cancer samples by taking advantage of partitioning around medoids based on informative low-dimensional features from the autoencoder. We constructed prediction models using 70% training sets and then predicted the labels of testing sets. To assess the robustness of the models, we adopted 10-fold cross-validation. Besides, we ran grid search cross-validation scheme to tune prediction algorithm parameters and sought the best model training set of parameters. To guarantee a more reliable performance evaluation, we repeated the random split, cross-validation, and grid search for all prediction models 10 times and took account of performance metrics on all evaluation experiments. We applied multiple preprocessing steps on both the training sets and testing sets. For gene expression, DNA methylation, and protein expression data, we adopted a robust scaling on the omics data using the means and the standard deviations. For miRNA expression and CNV data, we took advantage of unit-scale normalization [29].

### Evaluation metrics

To evaluate the prediction models comprehensively, we adopted multiple metrics, including accuracy, Kappa, balanced accuracy, ROC AUC. In addition, we adopted the brier score as the mean squared error between the expected probabilities for one specific subgroup and the predicted probabilities. The score ranges from 0 to 1, and a lower score suggests higher classification accuracy.

### Alternative PCA to the autoencoder

To show the robust performances of the autoencoder, we carried out principal components analysis (PCA) as the benchmark. We extracted the same number of principal components (100 PCs) as those low-dimensional features in the bottleneck of the autoencoder. We identified 15 survival-associated PCs using Cox-PH model and then we clustered the samples using the same partitioning around medoids for breast cancer patients.

### Ranking omics signatures associated with survival prognosis

Although the deep neural network is proved to be effective and robust in subgrouping breast cancers patients, the network is often perceived as a black box and it’s hard to derive informative biological insights from the model. To tackle the problem, we employed Olden’s algorithm to the neural network and extracted and ranked these potential multi-omics signatures according to their relative importance when subgrouping the samples [30]. Finally, we could deem those signatures ranking on the top and bottom as landmark multi-omics signatures contributing to the breast cancer patient stratification.

## Results

### Survival associated BRCA subtypes are identified

From the TCGA BRCA project, we obtained 501 patient samples that included gene expression, copy number variation (CNV), DNA methylation, protein RPPA expression, miRNA expression, and somatic mutation profiles. These multi-omics data were preprocessed as described in the “Methods” section, and finally, we obtained 18,596 gene expression features, 23,554 CNV features, 20,149 DNA methylation features, 186 protein expression features, 835 miRNA expression features, and 212,701 somatic mutation features in these patients. The workflow is shown in **Figure 1A**. After the model training and parameter optimization, we finally adopted the architecture setting (**Figure 1B** and **Supplemental Figure 1**). To note, early stopping terminated the model training and made the autoencoder avoid overfit as shown in **Figure 2B**. After that, we extracted 100 low-dimensional features from the 100 nodes of the bottleneck as new features (**Figure 2A**). To identify survival-associated low-dimensional features, we then built univariate CoX-PH model on each low-dimensional feature and obtained 16 features significantly associated with survival prognosis (wald and log-rank p-value < 0.05). Based on these 16 features, we performed partitioning around medoids in breast cancer patients, with cluster testing numbers from 2 to 5. By evaluating silhouette index and calinski-harabasz index, we determined that 2 clusters would be the best setting for clustering the patients (**Supplemental Figure 2A**). Interestingly, the 2 clusters classified the patients into 2 subgroups that represent two significantly different survival groups (Log-Rank P-value=0.0255), including low-risk subgroup S1 and high-risk subgroup S2 (**Figures 2C and 2D**). Considering S1 and S2 subgroups are informative for the clinical practice, we thus determined to adopt these two subgrouping criteria. In contrast, we found 3-clusters criterion, 4-clusters criterion, 5-clusters criterion for subgrouping breast cancer could not produce informative classification as shown in **Supplemental Figures 2C, 2D, and 2E**.

**Figure 1.**
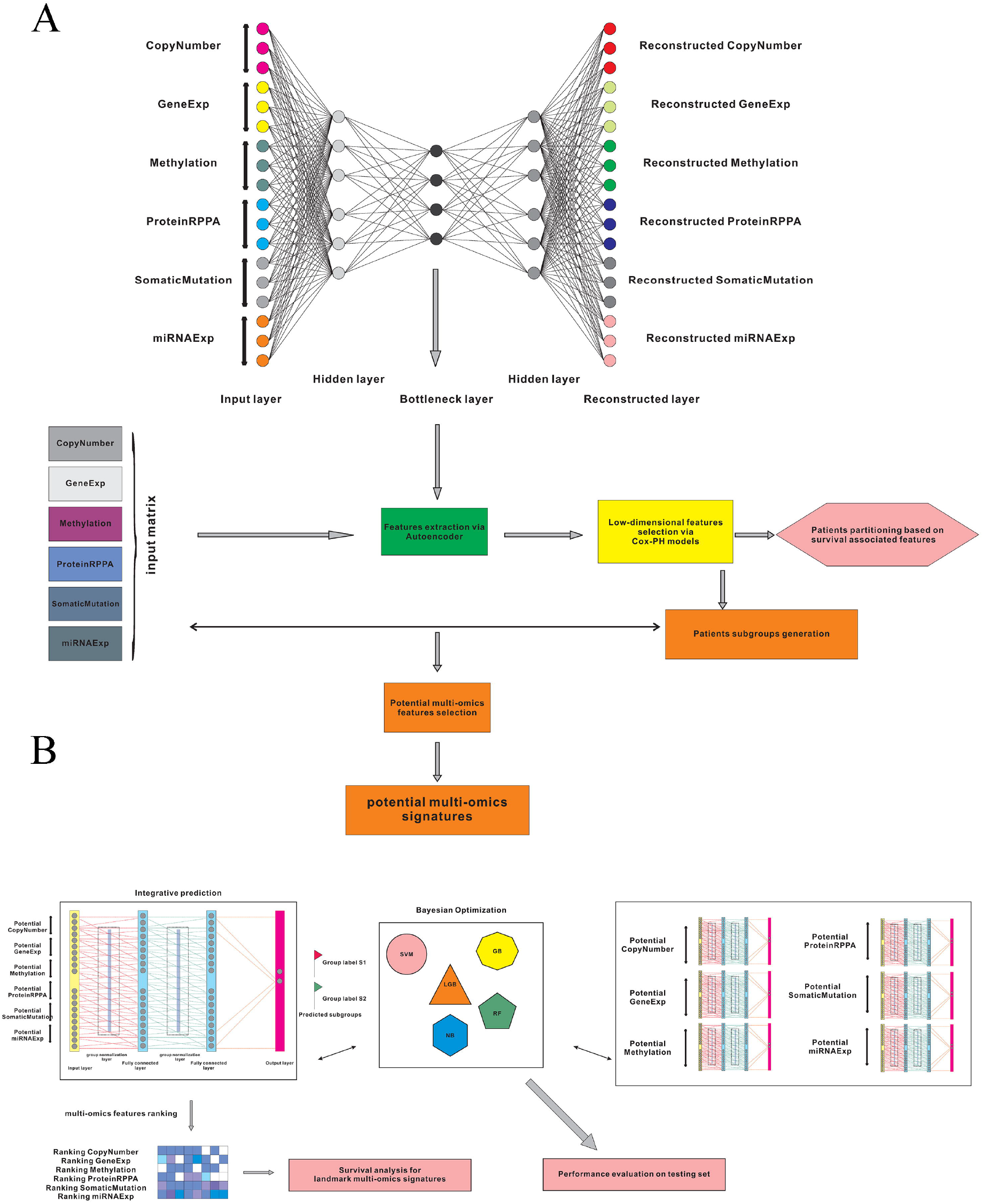
The workflow for this work. (A) Multi-omics data integration and patient stratification. (B) Prediction model training, performance evaluation, and multi-omics signature ranking.

**Figure 2.**
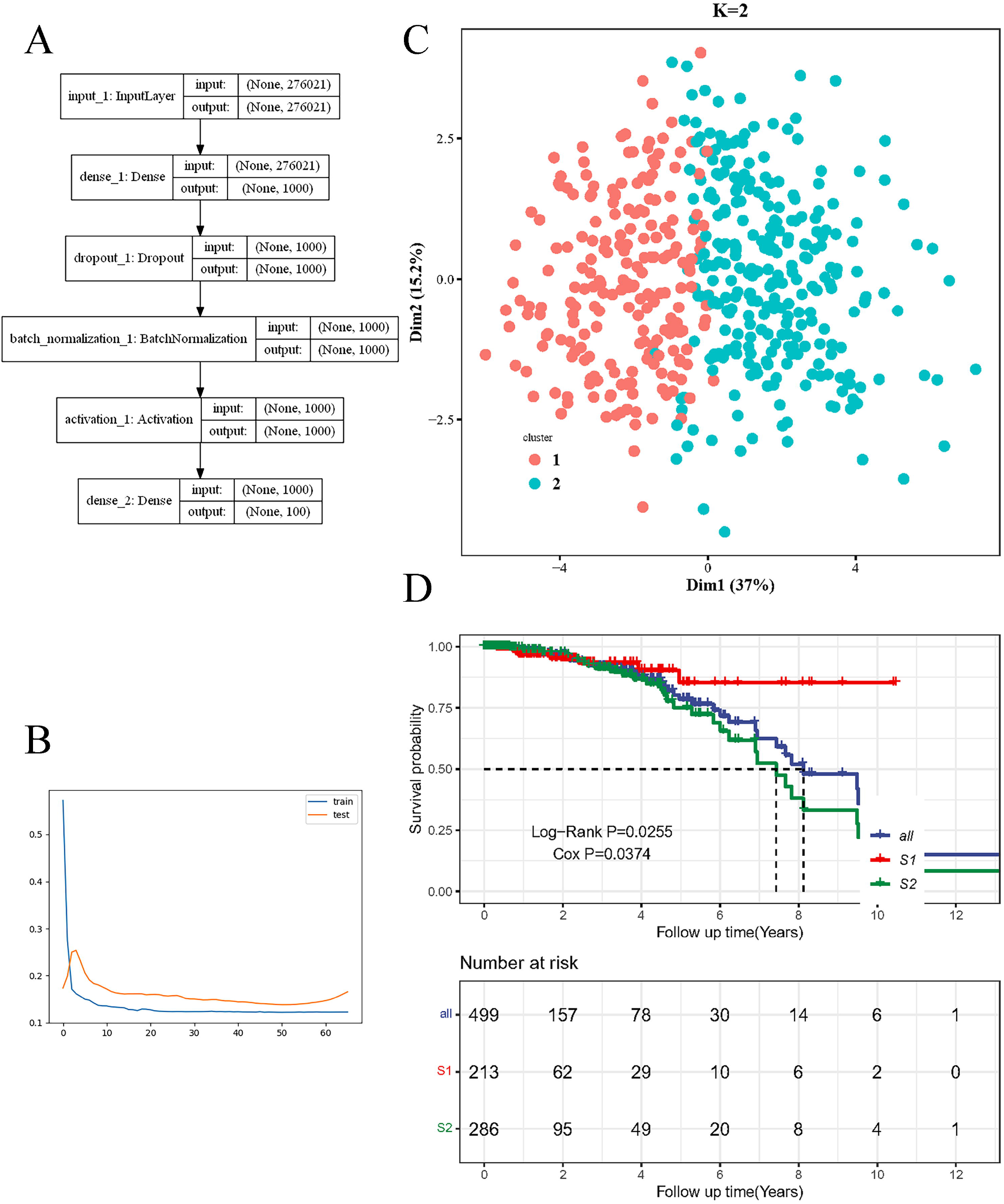
Deep learning model for breast cancer patient stratification. (A) The architecture of the encoder we established. (B) Training and test loss iteration during model optimization. (C) Breast cancer patient subgrouping visualization based on survival-associated low-dimensional features from the autoencoder. (D) Survival analysis for high-risk subgroup and low-risk subgroup based on survival-associated low-dimensional features from the autoencoder.

To corroborate superiority from our optimized autoencoder, we also carried out principal component analysis (PCA) on breast cancer patients. After picking up the top 100 PCs, we filtered and got 15 survival-associated PCs using Cox-PH model. Using partitioning around medoids, we found that these PCs failed to distinguish two subgroups from each other well (**Supplemental Figure 2B**).

### Differential multi-omics profiles characterize high risk and low-risk subgroups

Based on the two survival associated subgroups generated by partitioning around medoids, we identified 2,122 potential CNV features, 12,658 potential gene expression features, 6,419 potential methylation features, 38 potential protein expression features, 121 potential somatic mutation features, and 417 potential miRNA expression features that may contribute to the difference between the high-risk subgroup and low-risk subgroup (Wilcoxon test, P-value<=0.05). To enable the discovery of potential therapeutic targets in high-risk breast cancer, we selected top multi-omics features as putative multi-omics signatures, including 424 CNV signatures, 1,265 gene expression signatures, 641 methylation signatures, 38 protein expression signatures, 121 somatic mutation signatures, and 417 miRNA signatures.

As shown in **Figure 3A**, high-risk subgroup S2 harbored more *PEX2* copy number variation than low-risk subgroup S1 (Wilcoxon test, P-value=0.00084). *PEX2* was already predicted as a breast cancer risk locus from GWAS studies, our analysis indicated that increased CNV of *PEX2* was likely to be associated with breast cancer progression and development, rather than aberrant gene expression, methylation, protein expression, or somatic mutations of *PEX2* [31]. In addition, we found that decreased *STK11* gene expression (**Figure 3B**; Wilcoxon test, P-value<2.2e-16), decreased *CDH11* methylation (**Figure 3C**; Wilcoxon test, P-value=1.5e-6), increased *MIR628* miRNA expression (**Figure 3D**; Wilcoxon test, P-value<2.2e-16), decreased PRKCB protein expression (**Figure 3E**; Wilcoxon test, P-value=5.6e-6), the frameshift insertion of *PCLO* (**Figure 3F**; Wilcoxon test, P-value=7.4e-7) might contribute to breast cancer progression. Remarkably, many of these differential omics genes were found to be possible risk factors in breast cancer based on previous reports [32–35]. This suggests that our findings not only get cross verified by previous research but also add novel insights or possible mechanisms on how these risk genes may function in breast cancer progression. For example, it was reported that the elevated *CDH11* expression is associated with breast cancer metastasis. Combined with our finding above, one could infer reasonably the interaction between increased *CDH11* expression and decreased *CDH11* methylation was likely to contribute to high-risk breast cancer [33]. In addition, there were plenty of potential multi-omics signatures contributing to the difference between high-risk subgroup S2 and low-risk subgroup S1, including increased *HNF4G* CNV (Wilcoxon test, P-value=0.00057), decreased *USE1* gene expression (Wilcoxon test, P-value<2.2e-16), decreased *ESR1* methylation (Wilcoxon test, P-value=2e-7), increased *MIR874* miRNA expression (Wilcoxon test, P-value<2.2e-16), increased BECN1 protein expression (Wilcoxon test, P-value=0.0044), and increased translation start site SNP of *CEP290* (Wilcoxon test, P-value=2.8e-6) in high-risk S2 subgroup (**Supplemental Figures 3A, 3B, 3C, 3D, 3E, and 3F**). Again, many of them had supporting evidence to be associated with breast cancer in other ways [36–40], and these research works further corroborate our multi-omics findings and provide inspiration for breast cancer functional studies further.

**Figure 3.**
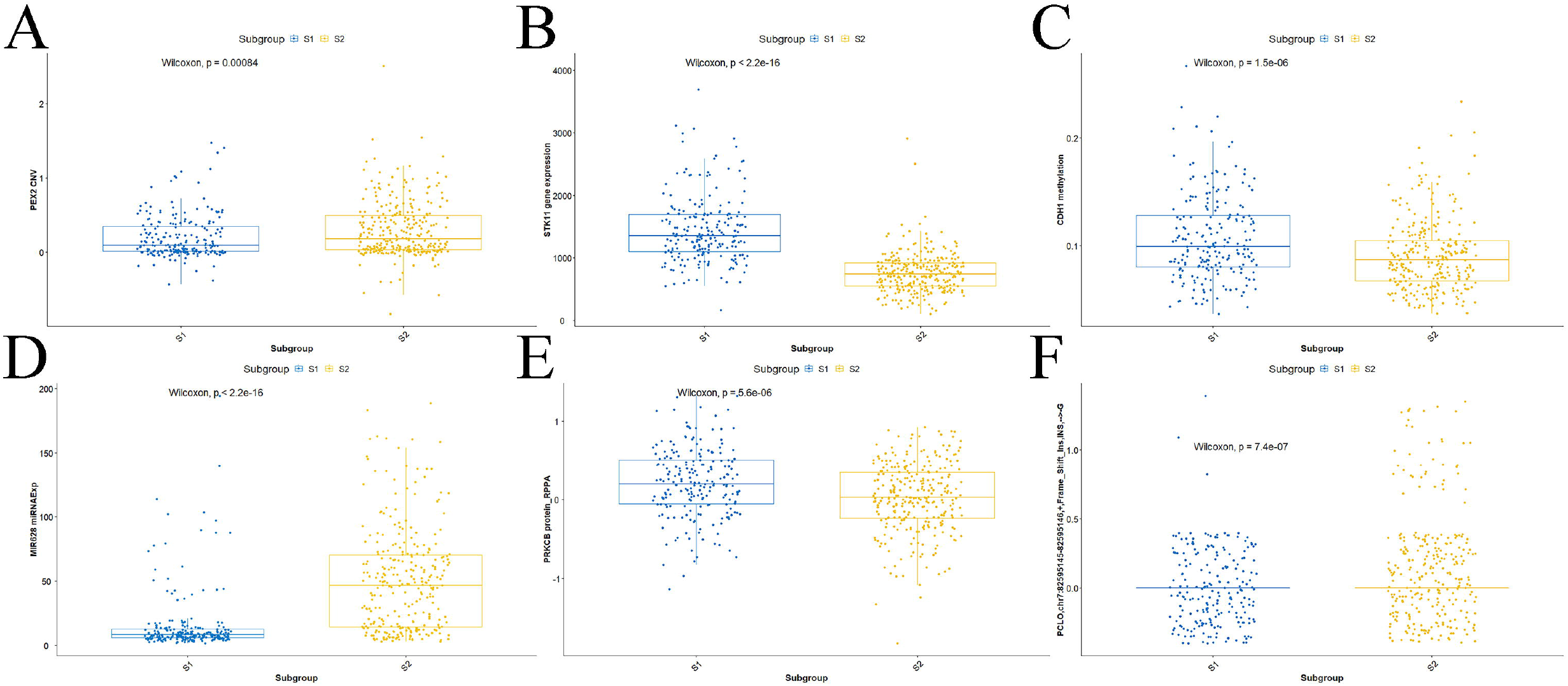
Profiles for multi-omics landmark signatures between high risk subgroup and low risk subgroup. (A) *PEX2* CNV (Wilcoxon test, P-value=0.00084). (B) *STK11* gene expression (Wilcoxon test, P-value<2.2e-16). (C) *CDH1* methylation (Wilcoxon test, P-value=1.5e-6). (D) *MIR628* miRNA expression (Wilcoxon test, P-value<2.2e-16). (E) *PRKCB* protein expression (Wilcoxon test, P-value=5.6e-6). (F) *PCLO* frame shift insertion (Wilcoxon test, P-value=7.4e-7).

*ATM, BARD1, BRCA1, BRCA2, MRE11A, NBN, PALB2, PTEN, RAD50, RAD51C, RAD51D,* and *XRCC2* are among the most critical risk factors contributing to triplenegative breast cancer as reported [41–53]. Interestingly, 11 of 12 these risk factors were also identified in our model as key multi-omics features distinguishing high-risk subgroup and low-risk subgroup, corroborating the validity of our subgrouping labels. We found three types of omics signatures of these risk genes contributing to the survival prognosis of breast cancer: the higher gene expression of *ATM, BARD1, BRCA1, BRCA2, NBN, PALB2, PTEN, RAD50, RAD51D,* and *XRCC2* was observed in high-risk subgroup S2; the higher CNV of *NBN* and *RAD51C,* the lower CNV of *XRCC2* was enriched in subgroup S2; as for DNA methylation, the lower methylation of *NBN, RAD50*, and *RAD51D,* the higher methylation of *BRCA2* was found frequently in subgroup S2 as well (**Supplemental Table S1**).

### Fully connected neural networks achieve robust predictive performance for classifying breast cancer patients based on multi-omics or single-omics data

To explore the value of the multi-omics signatures in subgrouping and predicting survival prognosis for breast cancer patients, here we developed and optimized multiple specified fully connected neural (FCN) networks customized for classifying breast cancer patients. We split the samples into 10 bins randomly using a 70/30 ratio where 70% training sets are for cross-validation and 30% testing sets are for testing. To exploit the potential performance, we carried out grid search for optimizing parameters for each neural network. Besides, we built multiple traditional machine learning as the benchmark, optimized and evaluated the predictive performance among these models, including support vector machine (SVM), random forest (RF), LogitBoost (LGB), and Naive Bayes (NB). To evaluate the models on testing sets comprehensively, we adopted multiple measures metrics, including accuracy, Kappa, balanced accuracy, specificity, sensitivity, ROC AUC, and Brier score.

The optimized FCN achieved excellent predictive performance when classifying breast cancer patients into two subgroups based on potential multi-omics signatures. Compared with other prediction models, FCN obtained better performance in these two valuable and practical evaluation metrics: for balanced accuracy, FCN achieved 93.34%, outperforming 89.01% of LGB, 84.94% of NB, 90.15% of RF, 86.70% of SVM; for Kappa, FCN achieved 86.21%, outperforming 79.05% of LGB, 70.41% of NB, 82.13% of RF, 72.32% of SVM (**Figure 4A** and **Supplemental Table S2**). In addition, FCN gained promising predictive performance in terms of accuracy, specificity, and sensitivity as well, suggesting 2-subgrouping for breast cancer patients based on multi-omics signatures is effective (**Figure 4A** and **Supplemental Table S2**).

**Figure 4.**
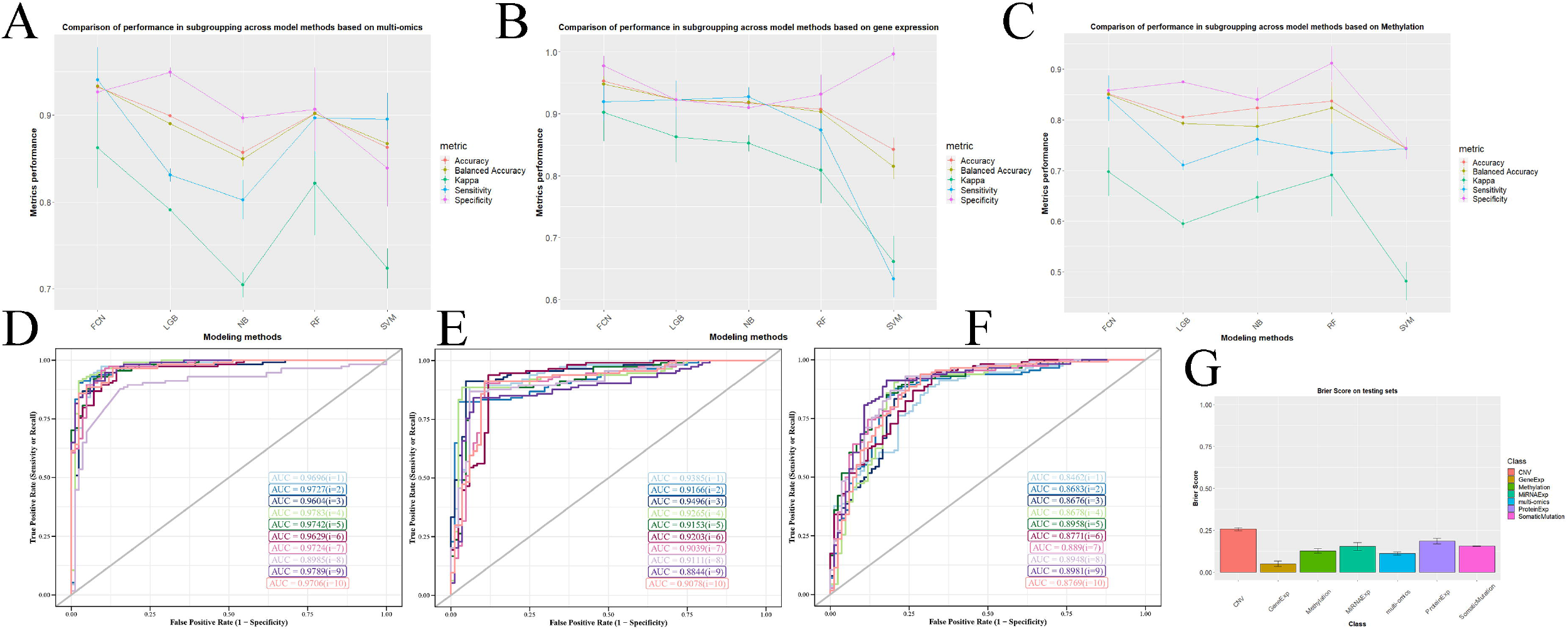
Performance evaluation for deep learning predictive models. (A) Comparison of performance in subgrouping across model methods based on multi-omics signatures in terms of accuracy, balanced accuracy, kappa, sensitivity, and specificity. (B) Comparison of performance in subgrouping across model methods based on gene expression signatures in terms of accuracy, balanced accuracy, kappa, sensitivity, and specificity. (C) Comparison of performance in subgrouping across model methods based on methylation signatures in terms of accuracy, balanced accuracy, kappa, sensitivity, and specificity. (D) Comparison of performance in subgrouping across model methods based on multi-omics signatures in terms of ROC AUC. (E) Comparison of performance in subgrouping across model methods based on gene expression signatures in terms of ROC AUC. (F) Comparison of performance in subgrouping across model methods based on methylation signatures in terms of ROC AUC. (G) Prediction performance in subgrouping for FCNs across multi-omics signatures and single-omics signatures in terms of brier score.

Besides, we explored the scenarios where there were limited available omics signatures. Interestingly, FCN still performed well in these situations and performed better than other models. When it comes to single-gene expression signatures, FCN gained an excellent predictive performance: for balanced accuracy, FCN achieved 94.81%, outperforming 92.26% of LGB, 91.87% of NB, 90.27% of RF, 81.49% of SVM; for Kappa, FCN achieved 90.22%, outperforming 86.30% of LGB, 85.25% of NB, 80.88% of RF, 66.13% of SVM (**Figure 4B** and **Supplemental Table S2**). Another robust example is single-methylation signatures where FCN performed better than other models: for balanced accuracy, FCN achieved 85.04%, outperforming 79.27% of LGB, 78.67% of NB, 82.32% of RF, 74.34% of SVM; for Kappa, FCN achieved 69.72%, outperforming 59.49% of LGB, 64.72% of NB, 69.12% of RF, 48.11% of SVM (**Figure 4C** and **Supplemental Table S2**). We could get similar trends when we evaluated these models in terms of accuracy, specificity, and sensitivity and the scenario of other single-omics signatures, including single-CNV, single-protein expression, single-somatic mutation, single-miRNA expression, suggesting 2-subgrouping for breast cancer patients based on single-omics signatures (or incomplete multi-omics) is feasible (**Supplemental Figures 4A, 4B, 4C and 4D**). For a comprehensive performance comparison based on multi-mocs and these six single-omics signatures in these five metrics, please refer to **Supplemental Table S2**.

Furthermore, we evaluated the optimized FCNs in the aspect of ROC AUC and Brier scores, and these FCNs were proved to be robust. For multi-omics signatures, FCN achieved 0.964±0.024 AUC and 0.112±0.010 Brier score, suggesting the complex functions in the fully connected neural network could capture the intrinsic relationship between multi-omics signatures and subgroup labels successfully (**Figures 4D and 4G**). Besides, FCNs based on single-omics signatures showed good performance. For example, FCN based on single-gene expression signatures achieved 0.917±0.018 AUC and 0.049 ± 0.016 Brier score and FCN based on single-methylation achieved 0.878 ±0.016 AUC and 0.126 ±0.013 Brier score (**Figures 4E, 4F and 4G**). We observed similar trends when we evaluated FCNs in the situation of other single-omics signatures (**Figure 4G**, **Supplemental Figures 5A, 5B, 5C, and 5D**). All these results suggest that these potential multi-omics signatures can provide promising values and the optimized FCNs manage to capture the intrinsic relationship between these omics signatures and the survival prognosis of breast cancer patients. For a comprehensive performance comparison based on multi-mocs and these six single-omics signatures in these two metrics, please refer to **Supplemental Table S3 and Table S4**.

### Ranking of landmark omics signatures unveils potential therapeutic targets

Although the optimized FCNs show excellent performance in classifying breast cancer patients into high-risk and low-risk subgroups, it’s hard to derive informative clues from the black-box models [54]. Here we employed Olden’s algorithm to the optimized FCNs we trained from the multi-omics data and extracted importance scores for each potential omics signature [30].

The individual signature of protein expression contributes most to classifying breast cancer patients into risk-associated subgroups, suggesting aberrant protein expression is more likely to be associated with the survival prognosis of breast cancer directly (**Figure 5A and 5E**). To note, although there are 417 potential miRNA expression signatures, outweighing the number of potential protein expression signatures and potential somatic mutation signatures, these potential miRNA expression signatures contribute less to subgroup breast cancer patients than other potential single-omics signatures overall (**Figure 5E**).

**Figure 5.**
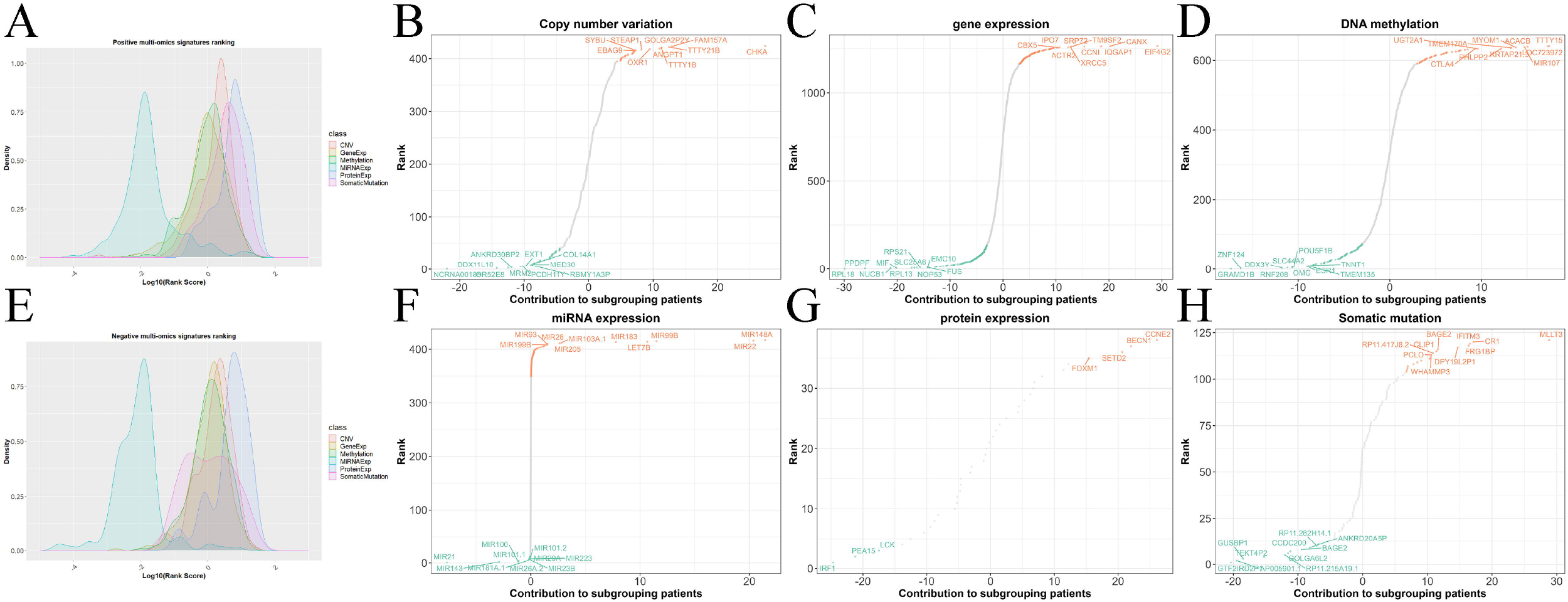
Ranking scheme of multi-omics signatures. (A) Ranking of positive multi-omics signatures. (B) Ranking of copy number variation signatures. (C) Ranking of gene expression signatures. (D) Ranking of DNA methylation signatures. (E) Ranking of negative multi-omics signatures. (F) Ranking of miRNA expression signatures. (G) Ranking of protein expression signatures. (H) Ranking of somatic mutation signatures.

For identifying potential therapeutic omics signatures, we ranked multi-omics signatures based on their respective omics layer. In the CNV signature layer, the top-ranked CNV signatures included *CHKA, FAM157A, TTTY21B, TTTY1B, GOLGA2P2Y, ANGPT1, STEAP1, OXR1, EBAG9, SYBU, NCRNA00185, DDX11L10, OR52E8, ANKRD30BP2, MRM2, PCDH11Y, EXT1, RBMY1A3P, MED30,* and *COL14A1* (**Figure 5B**). In the gene expression signature layer, the top-ranked gene expression signatures included *EIF4G2, CANX, IQGAP1, TM9SF2, CCNI, SRP72, XRCC5, IPO7, ACTR2, CBX5, RPL18, PPDPF, NUCB1, MIF, RPL13, RPS21, SLC25A6, NOP53, EMC10,* and *FUS* (**Figure 5C**). In the DNA methylation signature layer, the top-ranked methylation signatures included *TTTY15, LOC723972, ACACB, MIR107, MYOM1, TMEM170A, KRTAP21.3, PHLPP2, UGT2A1, CTLA4, GRAMD1B, ZNF124, DDX3Y, RNF208, OMG, SLC44A2, POU5F1B, ESR1, TMEM135,* and *TNNT1* (**Figure 5D**). In the miRNA expression signature layer, the top-ranked miRNA expression signatures included *MIR148A, MIR22, MIR99B, LET7B, MIR183, MIR103A.1, MIR205, MIR28, MIR93, MIR199B, MIR21, MIR143, MIR181A.1, MIR100, MIR101.1, MIR26A.2, MIR101.2, MIR29A, MIR23B,* and *MIR223* (**Figure 5F**). In the protein expression signature layer, the top-ranked protein expression signatures included CCNE2, BECN1, SETD2, FOXM1, TSC2, IRF1, PEA15, LCK, PXN, and PRKCB (**Figure 5G**). In the somatic mutation signature layer, the topranked somatic mutation signatures included *MLLT3* (chr9:20414343-20414343,+,Silent,SNP,AA>AG), *CR1* (chr1:207787753-207787753,+,Nonsense_Mutation,SNP,CC>CT), *IFITM3* (chr11: 320649-320649,+,Silent,SNP,GG>GA), *FRG1BP* (chr20:29632643-29632643,+,Missense_Mutation,SNP,TT>TC), *DPY19L2P1* (chr7:35131480-35131480,+,RNA,SNP,AA>AT), *GTF2IRD2P1* (chr7:72664015-72664016,+,RNA,INS,-- >-G), *AP005901.1* (chr18:15317007-15317007,+,RNA,DEL,CC>C-), *GUSBP1* (chr5:21490969-21490970,+,RNA,DEL,TTTT>TT-), *TEKT4P2* (chr21:9907668-9907668,+,RNA,SNP,GG>GA), *RP11-215A19.1* (chr4:187284501-187284501,+,RNA,DEL,GG>G-) (**Figure 5H**). To note, a significant portion of these topranked top signatures we identified were already found to be involved in breast cancer signaling in various biological pathways or mechanisms [55–69]. For a comprehensive ranking of landmark omics signatures in patient classification, please refer to **Supplemental Table S5**.

### Putative therapeutic targets identified from multi-omics deep learning

Although there were more than 3,000 potential multi-omics signatures in our result, the top-ranked signatures were thought to be more informative and valuable. We selected the top 60 landmark multi-omics signatures, and these included 22 gene expression signatures, 7 CNV signatures, 2 miRNA expression signatures, 9 methylation signatures, 10 protein expression signatures, and 10 somatic mutation signatures.

We performed univariate Cox-PH models using each of the top 60 omics signatures, and identified 16 survival-associated multi-omics signatures out of these signatures, including 11 gene expressions signatures, 2 miRNA expression signatures, 1 protein expression signature, 1 somatic mutation signature, and 1 CNV signature (**Figure 6A**). For example, we found that low gene expression of *ACTR2* (**Figure 6B**, Log-Rank P-value=0.0106), nonsense mutation (chr1:207787753-207787753,+,Nonsense_Mutation,SNP,CC>CT) in *CR1* (**Figure 6C**, Log-Rank P-value =0.00263), low protein expression of LCK (**Figure 6D**, Log-Rank P-value =0.00263), high miRNA expression of *MIR22* (**Figure 6E**, Log-Rank P-value P=0.0254), high CNV of *FAM157* (**Figure 6F**, Log-Rank P-value P=0.04), low expression of *RPL13* (**Figure 6G**, Log-Rank P-value P=0.016) were all associated with significantly poor breast cancer prognosis.

**Figure 6.**
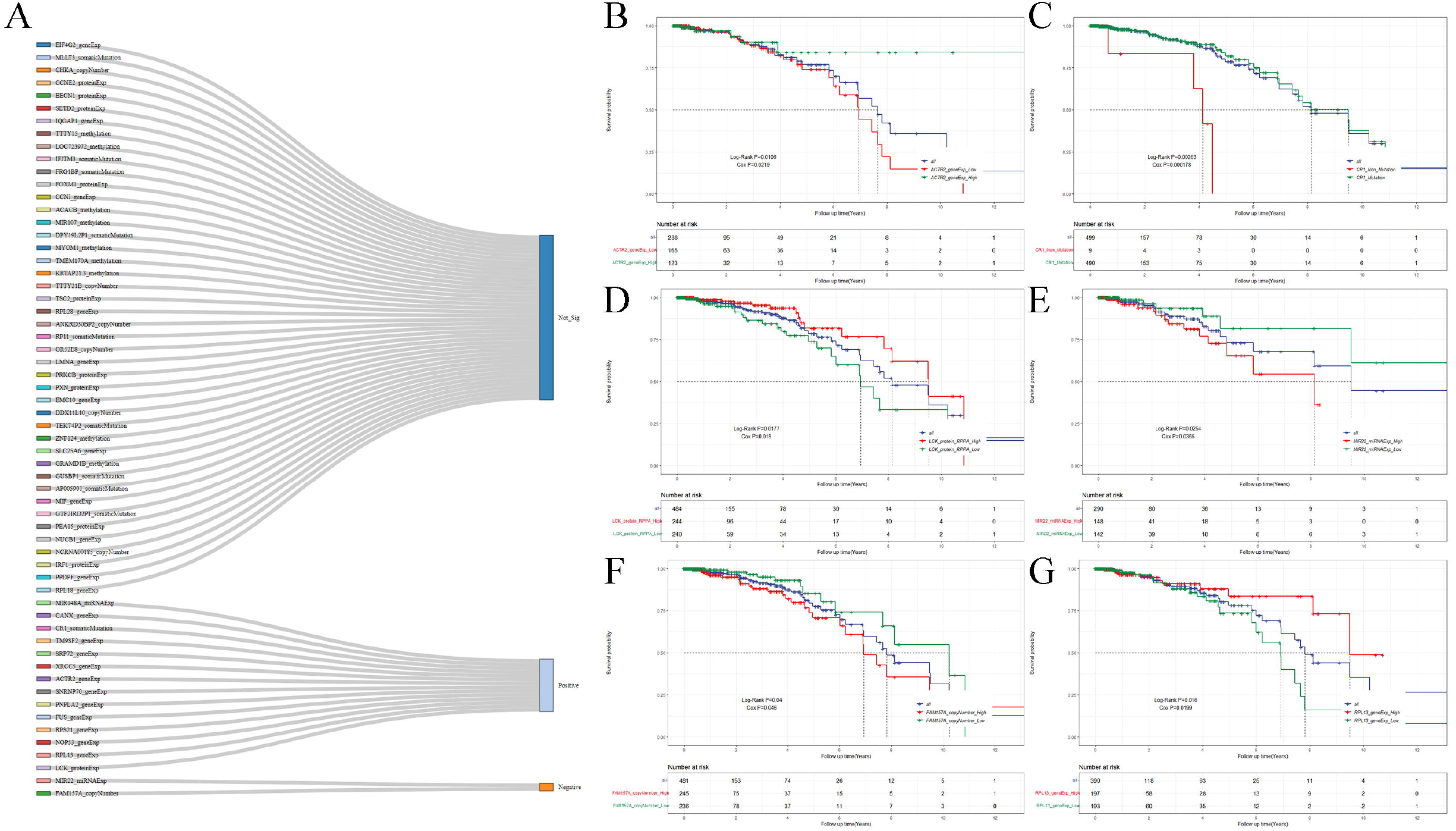
Multi-omics based identification of potential therapeutic targets for breast cancer. (A) Top 60 potential therapeutic multi-omics target candidates. (B) *ACTR2* gene expression (Log-Rank P-value=0.0106). (C) *CR1* mutation (Log-Rank P-value=0.00263). (D) LCK protein expression (Log-Rank P-value=0.0177). (E) *MIR22* miRNA expression (Log-Rank P-value=0.0254). (F) *FAM157* CNV (Log-Rank P-value=0.04). (G) *RPL13* gene expression (Log-Rank P-value=0.016).

We identified putative therapeutic targets for high-risk breast cancer, (top ranking) including low gene expression of *CANX* (Log-Rank P-value=0.0164), *FUS* (Log-Rank P-value=0.0379), *NOP53* (Log-Rank P-value=0.00128), *RPS21* (Log-Rank P-value=0.00424), *SNRNP70* (Log-Rank P-value=0.0154), *SRP72* (Log-Rank P-value=0.00706), *TM9SF2* (Log-Rank P-value=0.0327), *PNPLA2* (Log-Rank P-value=0.00069), *XRCC5* (Log-Rank P-value=0.0368), and low miRNA expression of *MIR148A* (Log-Rank P-value=0.0217) (**Supplemental Figures 6, 7 and 8**). Reassuringly, most of the top-ranked targets we identified were previously proved to be involved in breast cancer progression except *FAM157* [55–69]. For example, it was reported that the actin-related protein ARP2 encoded by *ACTR2* is the main component of Arp2/3 protein that promotes breast cancer proliferation and metastasis [55]. Another example is that the expression of cancer-specific *CR1* is decreased in triple-negative breast cancer [56]. Together, all these results further corroborate our multi-omics findings and may gain new insights into breast cancer development and accelerate precision drug discovery in the future.

## Discussion

Heterogeneity is one of the main limitations for understanding and diagnosing breast cancer. Effective subgrouping method to classify breast cancer patients into high-risk and low-risk subgroups is promising in clinical practice in the field. To our knowledge, we are the first to use the deep learning framework to integrate comprehensive multi-omics, including gene expression, miRNA expression, CNV, protein expression, somatic mutations, and methylation in breast cancer. Our high-risk and low-risk subgrouping strategy will benefit the diagnosis of breast cancer and provide a survival prognosis guideline for patients.

The optimized FCN models from multi-omics data achieve robust predictive performance for classifying breast cancer patients not only on multi-omics but also single-omics data, making our deep learning framework more precise and feasible in the reality. Based on the prediction model, we derived plenty of interesting biomarkers associated breast cancer risk and progression. For example, *CDH11* is already found to be a risk factor in breast cancer, however, the mechanism of how *CDH11* may play in breast cancer is still unknown [33]. We discovered inhibitive *CDH11* methylation will contribute to poor prognosis in breast cancer rather than proliferative *CDH11* methylation, aberrant *CDH11* gene expression, protein expression, somatic mutation, and CNV. Other biological results from our subgrouping will benefit the understanding of breast cancer in the multi-omics setting as well.

We evaluated and ranked the potential landmark multi-omics signatures and explored the importance of these signatures overall or from an individual perspective. Based on ranking of these multi-omics signatures, we discovered quite a few promising multi-omics signatures that are significant with survival prognosis and may serve as potential therapeutic targets to treating high-risk breast cancer in the future.

## Key Points

- A robust deep learning model to integrate multi-omics data in breast cancer systematically and explore breast cancer patient subtypes with survival prognosis.
- Accurate classification of breast cancer is critical with significant clinical implication, and our prediction models could robustly predict patient cancer risks even with incomplete omics data.
- Our deep learning model could identify landmark omics signatures associated with cancer development, and provide novel and effective therapeutic targets for hard-to-treat cancer subtypes.

## Supplementary Data

Supplementary data are available online.

## Conflicts of Interest

No conflicts of interest are disclosed by authors.

## Acknowledgements

The authors are grateful to contributions from TCGA Research Network Analysis Working Group, and also acknowledge the Biomedical Research Computing Facility at UT Austin, and Texas Advanced Computing Center (TACC) for assistance.

## Funding

This work was supported by the National Institutes of Health grants GM133658 (to S.Y.) and GM133406 (to N.S.), and Komen Foundation grant CCR19609287 (to S.Y.). N.S. is a CPRIT Scholar in Cancer Research with funding from the Cancer Prevention and Research Institute of Texas (CPRIT) New Investigator Grant RR160021.

## Notes

### Competing Interest Statement

The authors have declared no competing interest.

